# Selection shapes synonymous stop codon use in mammals

**DOI:** 10.1101/618157

**Authors:** Cathal Seoighe, Stephen J. Kiniry, Andrew Peters, Pavel V. Baranov, Haixuan Yang

**Affiliations:** School of Mathematics, Statistics and Applied Mathematics, National University of Ireland Galway, Galway, Ireland; School of Biochemistry and Cell Biology, University College Cork, Cork, Ireland

## Abstract

Phylogenetic models of the evolution of protein-coding sequences can provide insights into the selection pressures that have shaped them. In the application of these models synonymous nucleotide substitutions, which do not alter the encoded amino acid, are often assumed to have limited functional consequences and used as a proxy for the neutral rate of evolution. The ratio of nonsynonymous to synonymous substitution rates is then used to categorize the selective regime that applies to the protein (e.g. purifying selection, neutral evolution, diversifying selection). Here, we extend these models to explore the extent of purifying selection acting on substitutions between synonymous stop codons. Using a large collection of coding sequence alignments we estimate that a high proportion (approximately 57%) of mammalian genes are affected by selection acting on stop codon preference. This proportion varies substantially by codon, with UGA stop codons far more likely to be conserved. Genes with evidence of selection acting on synonymous stop codons have distinctive characteristics, compared to unconserved genes with the same stop codon, including enrichment for specific functional properties. Notably, genes with conserved stop codons have longer 3’ UTRs and are associated with shorter mRNA half-life than other genes. The coding regions of these genes are also much more likely to be under strong purifying selection pressure. Our results suggest that the preference for UGA stop codons found in many multicellular eukaryotes is selective rather than mutational in origin.

## Introduction

The standard genetic code includes three stop codons, UAG, UAA and UGA, that signal the end of translation. In eukaryotes termination of translation involves a heterodimer of two release factors, eRF1 and eRF3. A stop codon within the A site of the Ribosome is recognized by eRF1, inducing conformational changes in the ribosome via the GTPase activity of eRF3, ultimately resulting in the release of the polypeptide chain from the ribosome [1, 2]. Although all three stop codons can be recognized by eRF1, the efficiency of translation termination differs significantly between them, ranging from UAA (highest fidelity of translation termination) to UGA (lowest), with UAG being intermediate [2]. The altered conformation of the ribosome accommodates the nucleotide downstream of the stop codon (the +4 nucleotide) within the A site [3] and this nucleotide can also have a substantial impact on the efficiency of translation termination [2, 4]. Sequences further downstream as well as upstream of the stop codon can also affect readthrough efficiency and specific sequence contexts have been reported to result in readthrough efficiencies as high as 17% in human cells [5].

Consistent with the higher efficiency of translation termination associated with purines at the +4 position (i.e. the base immediately downstream of the stop codon), G is over-represented at this position in mammalian genes [6], particularly in highly expressed genes [7]. By contrast, despite having the lowest efficiency of translation termination among the three, UGA is the most common stop codon in many multicellular eukaryotes [8]. The relative frequencies of the three stop codons are dependent on multiple factors and strongly associated with regional variation in GC content [9]. The frequency of UAA is strongly negatively correlated with GC content, while the use of UAG and particularly of UGA increases with increasing GC content. This suggests a mutational origin for the variation in stop codon use, as mammalian GC content may be largely determined by mutational effects, reflecting variation in dNTP abundance across S-phase that favours the incorporation of A and T nucleotides in late-replicating DNA [10]. However, the relationship with GC content is less strong for stop codon use than for sense codons and this has been interpreted as evidence that selection also contributes to the choice of stop codon [9]. This is further supported by strong functional biases associated with stop codon use [9].

Although a preference for efficient translational termination could explain variation in stop codon preference between different gene classes, it does not explain why UGA, the least efficient stop codon, remains the most frequent stop codon in many multicellular eukaryotes, including human, in which it occurs in close to 50% of protein-coding genes. Programmed readthrough of the stop codon was discovered in viruses and enables their typically compact genomes to increase their coding capacity (reviewed in [11]). Only a limited number of cases of functional readthrough have been confirmed in higher eukaryotes; however, the application of genome-scale techniques has recently suggested over a hundred human genes as candidates for functional readthrough [12]. Different techniques have revealed candidates with relatively low overlap, suggesting either that some or all of the methods have a high false positive rate or that many more mammalian genes may be affected by functional readthrough. Stop codon readthrough may also have an important regulatory role, potentially involving mechanisms that degrade readthrough products [13]. Yordanova et al. [14] recently proposed an intriguing model whereby low-level readthrough of a stop codon may play a role in gene regulation and mRNA quality control by placing a constraint on the total translational capacity of an mRNA. Under this model, ribosomes that continue past the stop codon form a queue, backing up from a downstream ribosome stalling site. The rate of stop codon readthrough together with the length of the interval between the stop codon and the stall site control the number of times the mRNA can be translated before the ribosome queue backs up as far as the stop codon, inhibiting translation. Currently it is not known how widespread this mechanism is; however, if it is common it may have an impact on stop codon use, as the different readthrough efficiencies of the different stop codons would have implications for the number of times the mRNA is translated. Termination of translation is a slower process than amino acid incorporation, hence stop codons themselves are often used as ribosome stalling sites. Such stalling may affect mRNA stability via NoGo decay [15], but may have other important functions as in the above example or in the example of *XBP1* where it is required for its unusual cytoplasmic mRNA splicing [16]. Yet another regulatory event that involves stop codons is programmed ribosomal frameshifting [17] that often takes place at stop codons, e.g. +1 frameshifting in all three human antizyme genes (OAZ1, OAZ2 and OAZ3) takes place at stop codons and their identity is highly conserved. These, or as yet undiscovered functional implications of stop codon choice, may provide a selective explanation for the markedly high abundance of UGA stop codons in multicellular eukaryotes.

Here we set out to assess the extent of purifying selection affecting stop codon evolution in mammals. We extended models of codon sequence evolution that have previously been used to assess selection acting on coding sequences [18] to encompass the stop codon and fitted mixture models to estimate both the strength of purifying selection and the proportion of genes affected by purifying selection acting on the stop codon. Our model enabled us to assess the statistical evidence for selection for individual genes. Genes with conserved stop codons showed striking physical characteristics and were also enriched for certain gene functional classes. Stop codon conservation was more strongly associated with the selective constraints acting on the coding sequence than with the GC content of the gene, suggesting a selective, rather than a mutational origin for the variation in stop codon use with GC-content.

## Results

### Model

Standard codon models are characterised by a 61 × 61 transition rate matrix, **Q**, that gives the instantaneous rate of transition between each pair of sense codons [18–21]. Typically, these models assume that codons evolve through single nucleotide substitutions, according to a continuous-time Markov process, so that the instantaneous rate of transition between codons differing at more than one position is zero. Here we extend models of codon evolution by proposing a full 64 × 64 rate matrix that includes the three stop codons. As implemented here, we set the rate of substitutions between sense and stop codons to zero. Although the point at which translation terminates may shift (resulting from mutations between sense and stop codons), we consider only aligned sequences with the stop codons positionally homologous to the end of the alignment and assume correctness of the sequence alignment. The model can be modified easily to allow mutations between stop and sense codons by the addition of parameters corresponding to the rates of these substitutions. Note that the stop codon UAA is accessible by a single base mutation from both of the other stop codons (UAG and UGA); however, the instantaneous rate of transition between UAG and UGA is zero, as it requires two single nucleotide substitutions. We note also that, unlike standard codon models, the transition probability matrix obtained by exponentiating our rate matrix is not irreducible and does not, therefore, have a unique stationary distribution (a chain starting with a sense/stop codon will remain a sense/stop codon). Several forms have been proposed for the generator matrix of codon substitution models, differing mainly in how differences in codon usage are modeled. The model of Muse and Gaut [19] sets the rate of substitution from codon *i* to codon *j* (which differ at a single nucleotide position, *k*) to be proportional to 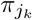, the equilibrium frequency of the nucleotide at position *k* of codon *j*. We follow this approach, as it has been found to be less prone to detecting spurious context-dependent effects than the version of Goldman and Yang [20], which sets the substitution rate to be proportional to the equilibrium frequency of codon *j* [22]. *The entries, q*_*ij*_, of the rate matrix of our model are therefore:

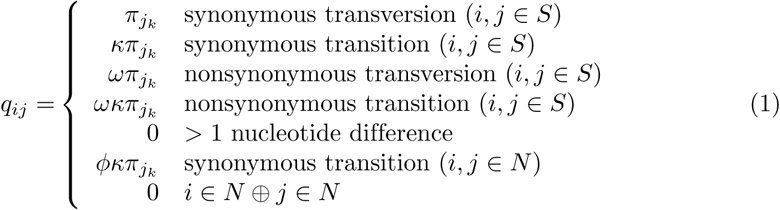

where *S* and *N* are the sets of sense and stop (or nonsense) codons, respectively (note that all mutations between stop codons are transitions). The last two conditions represent the extension of the model to accommodate stop codons. A parameter, *ϕ*, models the substitution rate between stop codons, relative to the rate of synonymous substitutions between sense codons. The last condition assigns a zero rate for substitutions between sense and nonsense codons. Although a parameter can easily be added to allow mutations between sense and nonsense codons (resulting in a shift in the stop codon position) as presented above the model consists of two subsets of intercommunicating states (corresponding to the sense and stop codons). This model could be fitted as a combination of a standard codon model and a 3×3 rate matrix for the stop codons; however, the model parameters would need to be estimated jointly and therefore we consider that this is best represented as a single reducible Markov process.

We implemented this model in R [23], optimizing parameters using the Nelder-Mead [24] method. We simulated data under the model and found that we could recover simulated values of *ϕ*, the parameter of interest, with no evident bias (Fig. S1a; slope = 1.008, estimated with robust regression, using an M estimator). We also simulated data under the Goldman and Yang [20] model, using empirical triplet frequencies obtained from intron sequences, to investigate if we could recover *ϕ*, when the simulated data were generated with a different version of the codon model to the one used for inference. We found that *phi* could still be recovered from the simulated data with little bias (Fig. S1b; robust linear model slope = 1.04; see Methods for details of the simulations).

### Proportion of stop codons under purifying selection

To estimate the proportion of stop codons evolving under the influence of purifying selection, we fitted our stop-extended codon model to the codon-aware alignments of mammalian orthologues, obtained from the OrthoMaM database [25], using a mixture distribution for the *ϕ* parameter. The mixture distribution consisted of two point masses, one with *ϕ* a free parameter (constrained to be < 1), corresponding to alignments with a stop codon evolving under purifying selection and another with *ϕ* fixed at 1, corresponding to neutral evolution – i.e. substitutions between stop codons occurring at a rate consistent with the rate of synonymous substitutions in the coding region. We then used maximum likelihood to estimate the two free parameters of this mixture model (the *ϕ* parameter for the constrained stop codons and the mixture weight parameter).

We estimated that 57% (Fig. 1) of mammalian stop codons are evolving under relatively strong (point mass for *ϕ* at 0.24) purifying selection. Using simulation we found that this estimate is not strongly dependent on modeling assumptions (Fig. S2). To investigate differences in selection pressure between genes with each of the three stop codons, we separated the genes into three groups, depending on which stop codon was found in the human gene. When we analysed these groups separately, we found that a substantially higher proportion of UGA stop codons are under selective constraint, compared to UAG and UAA stop codons. Bootstrapping over orthologue families suggests that the higher frequency of conservation in UGA stop codons is highly robust to sampling error (Fig. 1).

**Fig 1.**
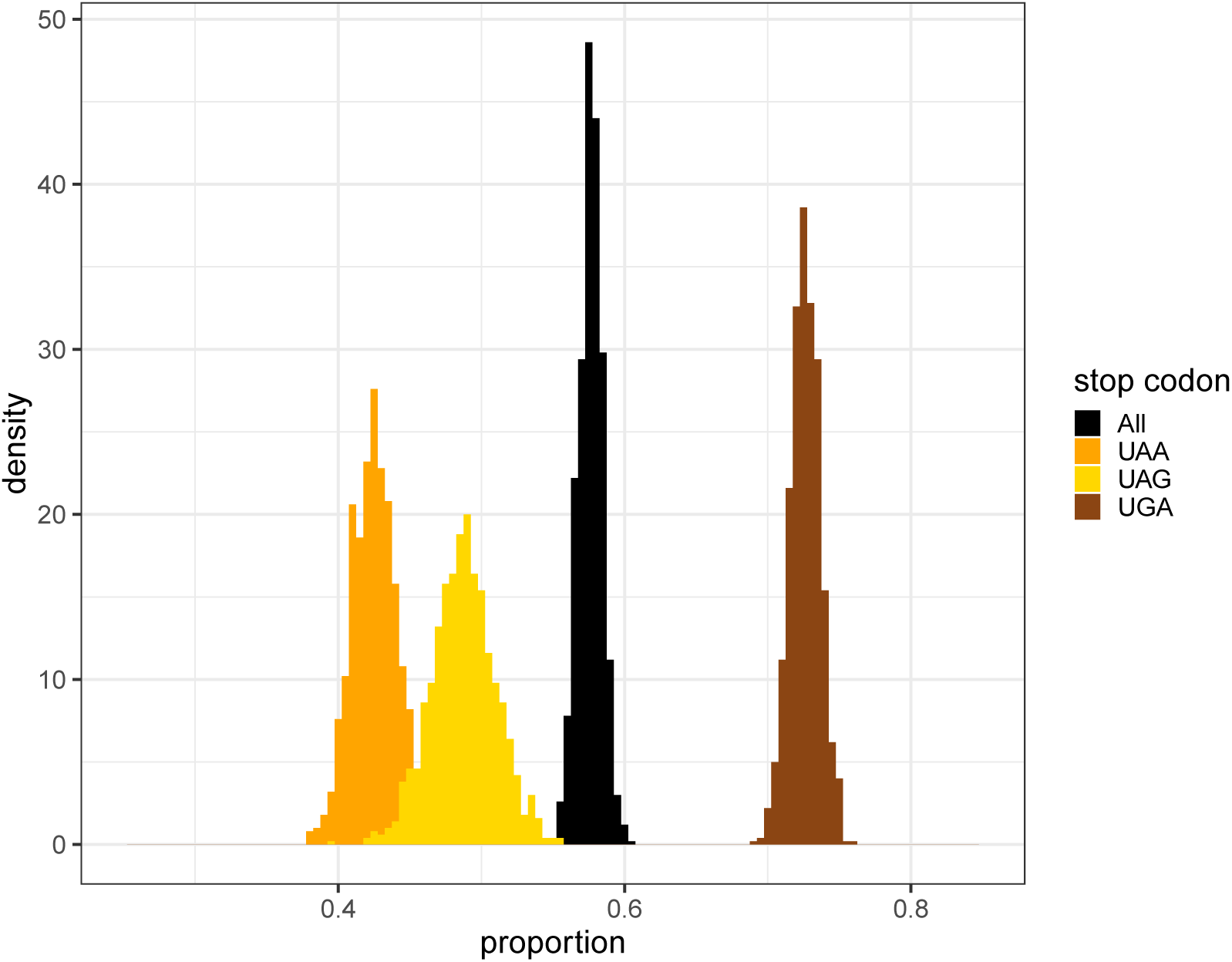
Mixture model results. Estimated proportions of stop codons under purifying selection for all genes (black) and for genes with UGA, UAG and UGA shown in brown, yellow and orange, respectively (or umber, amber and opal, according to the colour nomenclature for stop codons). Each histogram was derived from 1000 bootstrap replicates of the data.

### Identification of genes with conserved stop codon use

We also fitted our extended codon model to each orthologue family independently, estimating a separate value of *ϕ* for each gene. The rate of evolution of stop codons varied widely (Fig. S3). For 3,642 orthologue families (29.5% of those included in the analysis) the stop codon was completely conserved across the mammalian phylogeny, resulting in maximum likelihood estimates of zero for *ϕ*, while for some other genes the point estimate of *ϕ* was greater than one. An example of a gene in each category is provided in figure S4.

To assess the evidence for selection acting on stop codon use at the level of individual genes we used a likelihood ratio test, comparing the likelihood of the Null model with *ϕ* = 1 to the maximum likelihood of the alternative model with *ϕ* as a free parameter. For data simulated under the Null model the likelihood ratio test (LRT) statistic (twice the difference in the log likelihood between the null and alternative models) matched the expected *χ*^2^ distribution, with one degree of freedom (Fig. S5a). The fit with the *χ*^2^ distribution was less good when we simulated data using a model based on triplet nucleotide frequencies from introns (Fig. S5b; see Methods for details); however, even in this case the proportion of simulations for which the LRT statistic exceeded the critical value *and* for which *ϕ* was less than one was 0.058, approximately equal to the significance level (*α* = 0.05) applied. All genes for which the LRT statistic exceeded the critical value and for which the maximum likelihood estimate, 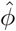, was less than one were considered as putatively under selective constraint. Using these criteria 27% of genes showed evidence of purifying selection acting on stop codon preference. We caution that some of these may be false positives, given the 5% significance threshold applied. The lower proportion (27%) of genes with conserved stop codons detected using the likelihood ratio test applied to individual genes, compared to the estimate from the mixture model, likely reflects limits of the power to reject the null hypothesis for individual alignments, as the null hypothesis was not rejected for some of the alignments with completely conserved stop codons. For 2,881 of the 3,642 genes with completely conserved stop codons, we reject the null hypothesis of neutral evolution (the null hypothesis was also rejected in favour of purifying selection for a further 447 genes with some stop codon substitutions). Failure to reject the null hypothesis even when the stop codon was completely conserved across the phylogeny was more likely to occur for UGA (neutral model rejected for 78% of completely conserved stop codons) or UAG (73%) stop codons than for UAA (95%) stop codons. The higher power for UAA results from the fact that UAA may mutate to UGA or UAG in a single point mutation, reducing the likelihood of passive conservation, relative to UGA and UAG, which can only mutate via single point mutations to UAA. Differences in power between stop codons means that the frequency of purifying selection cannot be compared between the stop codons on the basis of individual tests; however, this does not undermine the comparison based on the mixture model as described above, as the latter can accumulate evidence of frequent purifying selection across genes.

### Properties of genes with conserved stop codons

We compared the following properties between genes with conserved and non-conserved stop codons: 3’ UTR length,5’ UTR length, downstream open-reading frame length, coding sequence length, mRNA half-life, dN/dS, GC content, expression breadth, expression level. These properties were compared between all genes and between groups of genes, defined by the stop codon found in humans. Genes with putatively conserved stop codons had several striking features. Notably, they had longer 3’ untranslated regions (UTRs) than other genes (median of 1,183bp compared to a median of 877bp for the remainder of the genes in the dataset; p = 1 × 10^−39^). However, the lengths of 3’ and 5’ UTRs are strongly negatively correlated with GC content [26]. When we fitted a linear model to 3’ UTR length as a function of stop codon conservation and mRNA GC content we found the effect of conservation remained positive and significant (effect size estimate = 300bp; p = 7 × 10^−14^). Interestingly, the mRNA half-life of genes with conserved stop codons was shorter than that of other genes (Fig. 2). A strong effect was observed only for UGA stop codons with weak effects in opposite directions observed for UAG and UAA stop codons. For genes with a UGA stop codon (in human) half-life remained significantly associated with stop codon conservation when we fitted a logistic regression model with GC content, 3’ UTR length, CDS length and the ratio of nonsynonymous to synonymous substitution rates, *ω*, as predictor variables (p = 0.0004). Both the expression level and breadth of the stop-conserved genes were significantly higher than that of other genes (p = 1 × 10^−8^ and 2 × 10^−10^, respectively, using a Wilcoxon rank sum test). When these effects were investigated separately in genes with different stop codons (in human) they remained strongly statistically significant in genes with UGA and UAA stop codons but not in genes with UAG stop codons.

**Fig 2.**
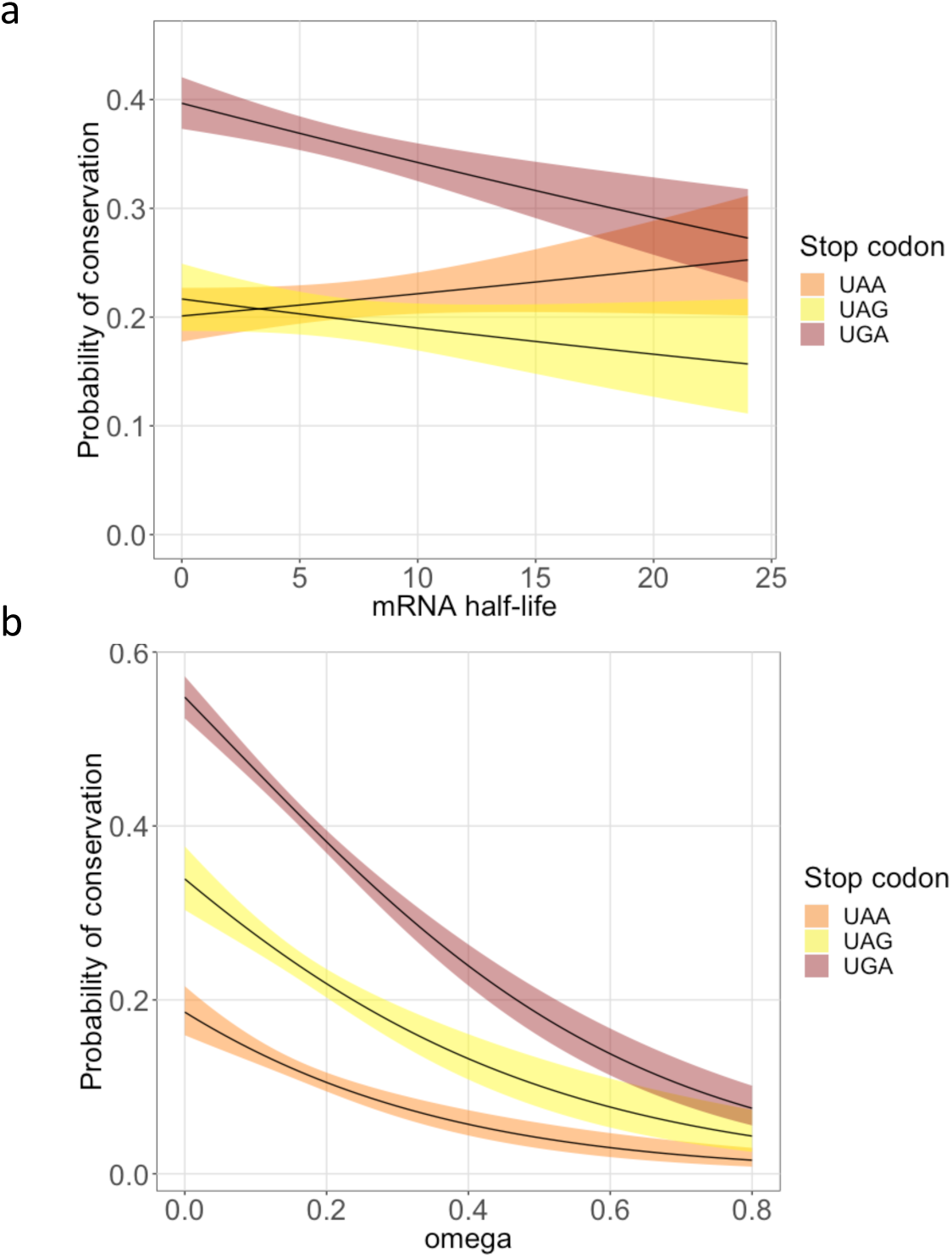
Relationship between stop codon conservation and mRNA half-life and coding sequence conservation. Estimated probability of stop codon conservation (and 95% confidence interval) as a function of (a) mRNA half-life and (b) *ω* for each stop codon type. Conservation is based on model comparison in (a) and on complete sequence conservation across the alignment in (b). Estimates are from a logistic regression model, which included the number of taxa for which the stop codon was positionally homologous with the end of the alignment as a covariate.

### Model-free analysis supports a major role for selection in shaping stop codon use

As an alternative to the model-based approach to defining conserved stop codons we investigated the characteristics of genes for which the stop codon was completely conserved across the entire alignment, regardless of whether there was sufficient evidence from the model to reject neutrality. We found that *ω*, the ratio of nonsynonymous to synonymous substitution rates, was strongly associated with stop codon conservation, with genes with low *ω* values (consistent with strong purifying selection acting on the CDS) having a much higher probability of having a stop codon that was completely conserved across the alignment (Fig. 2; this effect was also evident for the model based conserved set). We fitted a logistic regression model, treating complete conservation of the stop codon across the alignment as the outcome and with GC-content, stop codon (in human) and *ω* as predictors and including the interaction between stop codon and *ω* (i.e. allowing for different *ω* effects for different stop codons). Using the DynNom R package we provide an interactive interface to the statistical model at http://sonrai.nuigalway.ie:8001/CSeoighe. There was a striking effect of *ω* on the probability that a stop codon was conserved (Fig. 2). This suggests that conservation of the stop codon is influenced by selection, as genes with low values of *ω* are under strong selective constraint. Moreover, when we fitted an equivalent model but with GC-content and stop codon (in human) and included interaction terms, none of the interactions between stop codon and GC-content were significant, revealing no difference in the relationship with GC-content between the stop codons. Given the large variation in mutational patterns as a function of GC-content [10] this suggests that variation in mutation patterns is not the cause of differences in stop codon conservation. Further evidence that stop codon use is influenced by selection, rather than mutational effects comes from analysis of the frequency of each stop-associated triplet in the 3’ UTR. The frequency of UAG, UGA and UAA in the 3’ UTR in human (across all three forward frames) was 23%, 38% and 38%, respectively, suggesting that the excess of UGA over UAA stop codons is not mutational in origin, although the low abundance (22% in human) of UAG stop codons may be a mutational effect.

Characteristics of the set of genes with stop codons conserved across the mammalian alignment (the sequence-based set) were similar to those of the genes identified using the model (the model-based set) and, indeed, the majority (70%) of the genes that occurred in either group occurred in both. The sequence-based conserved gene set also had significantly longer 3’ UTRs. The mRNA half-life effect was more striking. In a logistic regression model with membership of this set as the outcome variable and including GC content, 3’ UTR length, CDS length, mRNA half-life and dN/dS ratio as predictors, the mRNA half-life coefficient was significantly different from zero for the complete set of genes (p = 1 × 10^−8^) as well as for the genes with UGA or UAG stop codons in human (p = 2 × 10^−6^ and 0.01, respectively), but not for genes with UAA stop codons in human (p = 0.68).

### Nonsynonymous but not synonymous divergence strongly predicts conservation of UGA stop codons

To investigate further whether conservation of the stop codon results from purifying selection or from a lower mutation rate or random chance we assessed the relationship between the probability of stop codon conservation and synonymous/nonsynonymous divergence. We obtained the number of nonsynonymous substitutions per nonsynonymous site (dN) and the number of synonymous substitutions per synonymous site (dS) for human-mouse orthologues from Ensembl [27]. Although, we already report above that the dN/dS ratio (i.e. *ω*) is predictive of stop codon conservation, these data allowed us to investigate the relationship with dN and dS separately, using values calculated independently of the alignments analyzed here. We found that dN was highly predictive of complete stop codon conservation but dS was only weakly predictive (Fig. 3). On average synonymous substitutions are under much weaker selection than nonsynonymous substitutions and dS is therefore much more reflective of the underlying mutation rate than is dN. Our observation that dS is a much weaker predictor of stop codon conservation than dN suggests that a lower mutation rate is not sufficient to explain conservation of the stop codon across mammals. It is possible that weak relationship between dS (human-mouse) and stop codon conservation is due to saturation of synonymous substitutions between human and mouse, given the relatively distant divergence of these species. Therefore, we also fitted the same model to complete sequence conservation as a function of divergence values with macaque, which diverged from humans much more recently. Again dN was strongly associated with stop codon conservation (at least for UGA and UAA stop codons), while dS was not (Fig. 3).

**Fig 3.**
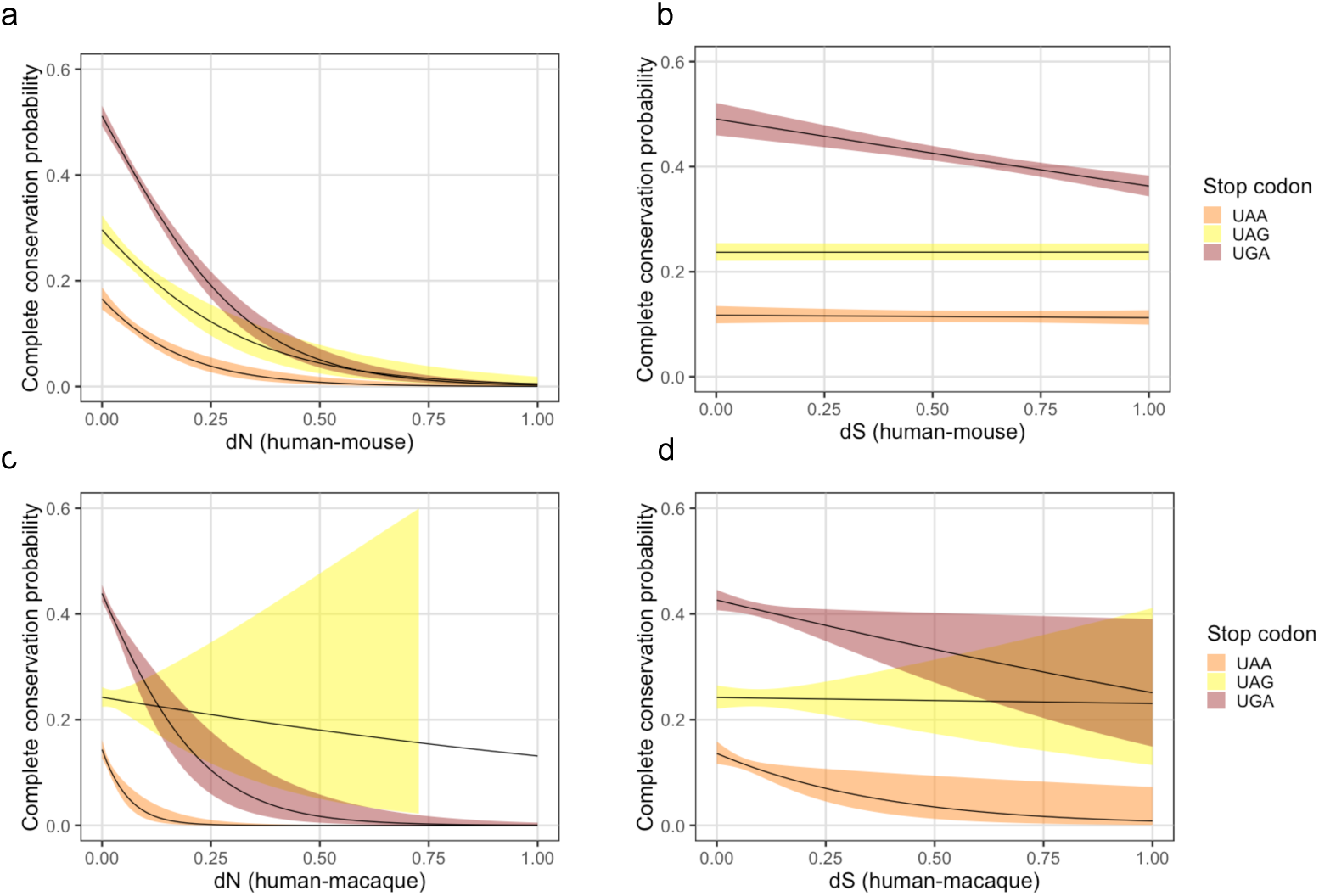
Stop codon conservation and nonsynonymous and synonymous distance. Estimated probability of complete sequence conservation (and 95% confidence interval) as a function of (a) dN between human and mouse (n = 12,266), (b) dS between human and mouse (n = 12,266), (c) dN between human and macaque (n = 12,083) and (d) dS between human and macaque (n = 12,083). In all cases the x-axis is truncated to 1, as most of the divergence values are lower than this. The number of taxa for which the stop codon was positionally homologous with the end of the alignment was included as a covariate in the model.

### Gene set analysis of genes with conserved stop codons

We performed gene set analysis on the genes with completely conserved stop codons using DAVID (version 6.8) [28, 29]. For this analysis we focussed on the set of genes for which the stop codon was completely conserved across the mammalian CDS alignment. We focused on this set, rather than the model-based set because the power to detect conservation using model comparison may differ between genes and systematic variation in detection power between genes can give rise to spurious results from gene set analysis [30]. The analysis was performed separately for each of the three stop codons and all genes with that stop codon (in human) were used as the background set. Sets of genes showing complete conservation of the UGA or UAA stop codon were both highly significantly enriched for genes involved in transcriptional regulation and, in particular, for homeobox transcription factors, including for HOX genes. There was very limited evidence of functional enrichment for genes with a UAG stop codon. Only a single gene set (Uniprot sequence feature, consisting of a polyserine region; adjusted p = 0.01) passed the 0.05 p-value threshold (adjusted using the Benjamini and Hochberg method).

## Discussion

We set out to determine the extent to which alternative stop codon use is affected by purifying selection in mammalian genes. By extending models that were developed to understand the selection pressures acting on amino acid encoding sequences, we estimated that the stop codon is subject to purifying selection in a large proportion (approximately 57%) of mammalian genes. The proportion under selection varies between the stop codons and is highest for genes with UGA stop codons (fig. 1). UGA is by far the most common stop codon in human and many other multicellular eukaryotes (close to 50% of human protein-coding genes have a UGA stop codon). Given the high rate of purifying selection affecting UGA stop codons we propose that the predominance of UGA codons is a result of selection rather than mutation. Stop codon use is strongly associated with GC-content [9] and large-scale variation in GC content across genomes has a mutational origin [10]. However, if the high abundance of UGA codons was mutational in origin we would expect that UGA codons in regions with high GC content would be much more likely to be conserved than UGA codons in low GC regions, given the strong relationship between GC content and stop codon use. This was not the case, as we found only a weak relationship between the probability of complete sequence conservation and GC content for all three stop codons.

Stop codon conservation may result from gene regulatory mechanisms that have a preference for the use of a specific stop codon. These mechanisms may include the control of translation capacity of mRNA molecules [14] and translational pausing, which has previously been associated with localization of an mRNA-ribosome-nascent chain [16]. Strong enrichment of UGA among conserved stop codons suggests that UGA may frequently be the preferred codon in such cases. In principle, it is possible that some cases of purifying selection acting on stop codons result from missed examples of UGA codons that encode selenocysteine, rather than signaling the end of translation. However, given the small number of genes encoding selenoproteins [31] and the large number of conserved UGA stop codons, this is very unlikely to explain a substantial proportion of the conservation we report.

The use of the synonymous substitution as a proxy for the rate of neutral evolution has been criticised, as it is known that synonymous substitutions may have functional consequences [32–36]. Codon models that include a variable synonymous substitution rate have been developed [37, 38]. By not incorporating variability in the synonymous substitution rate in the coding region we are effectively comparing the rate of synonymous stop codon substitutions to their expected rate, given the *mean* rate of synonymous substitution in the coding region. Given the extent of purifying selection acting on synonymous sites, the mean synonymous substitution rate is likely to be an underestimate of the neutral rate of evolution. However, because our objective here is to study purifying selection affecting synonymous stop codons underestimation of the neutral rate would make our method conservative (we would miss some genes under purifying selection, but the underestimate of the neutral rate should not cause false positives). We also observed many genes for which the maximum likelihood estimate of *ϕ* was greater than one (including *PARP1*, shown in fig. S4). It is possible that some of the genes with stop codons evolving at a rate greater than expected, given the synonymous rate, result from purifying selection acting on mutations between synonymous sense codons, but it is also possible that there is diversifying selection acting on stop codon use in some genes. However, the number of genes with *ϕ significantly* greater than one, was not more than we expected by chance (260 from a total of 12,336 genes at a significance threshold of *α* = 0.05). Using a very different method to ours, Belinky *et al.* [39] recently carried out an analysis of stop codon substitutions in a wide range of taxa. Although the majority of the taxa studied were microbial, they included three mammalian species. The authors reported an excess of substitutions to UGA stop codons, which they attributed to positive selection. However, consistent with our findings, they also report widespread purifying selection acting on synonymous stop codon mutations in primates, particularly for UGA [39].

Among the most striking properties of genes with conserved stop codon use was the decreased mRNA half-life for conserved genes with UGA stop codons (fig. 2). The length of the *3*’ UTRs is known to be inversely correlated with mRNA half-life and the conserved genes had longer 3’ UTRs; however, the reduced half-life in the UGA genes remained significant, even when we fitted a logistic regression model incorporating 3’ UTR length and GC-content as covariates. Although there may be many possible explanations for the lower half-life of these genes we note that the model proposed by Yordanova *et al.* [14] consisting of a mechanism to limit the number of times an mRNA molecule is translated may result in lowered half-life of the mRNAs affected because stalled ribosomes trigger mRNA decay [15]. Arriber*e et al.* found evidence of instability of proteins resulting from readthrough in *Caenorhabditis elegans* and human cells and noted that this has been reported to result in mRNA instability in the *HBA2* gene [13, 40]. Given the apparent ubiquity of protein instability resulting from readthrough, the higher readthrough rate for UGA codons and the shorter mRNA half-life of genes with conserved UGA stop codons, destabilization of mRNA, resulting from readthrough may be a common mechanism of controlling protein abundance. However, selection acting against synonymous mutations between stop codons may have reasons other than readthrough. In this regard we note a recent analysis of stop codon readthrough in *Saccharomyces cerevisiae* and *Drosophila melanogaster* ([41]) that suggests that many stop codon readthrough events may be molecular errors rather than having a specific function.

Previous studies have investigated stop codon sequence conservation, for example as one of the strands of evidence of functional translational readthrough [42–44]. Our study is distinct in that we do not set out to investigate a specific function of conserved stop codons but, instead, to estimate the frequency and strength of selection acting on synonymous stop codon use and to investigate the properties of the associated genes, in general. In principle, inference of stop codon conservation using our extended codon model is preferable to inference based on sequence conservation alone, as the latter does not take into account differences in GC-content and mutation rates between genomic regions. Our method explicitly models variation in codon usage, through incorporation of parameters (estimated empirically from the entire CDS alignment) for codon equilibrium frequencies. However, the total tree length and number of taxa in the mammalian orthologue alignments from OrthoMaM [25] was only just sufficient in many cases and in some other cases insufficient to reject the neutral model, even in the presence of complete conservation of the stop codon across all taxa. As the size of the families of reliably aligned coding sequences increases, the power to estimate accurately the strength of purifying selection acting on synonymous stop codons will increase. This will allow much more accurate identification of the affected genes and represents an example of the value of continued efforts to sequence the genomes of an ever increasing range of organisms and of the curation and alignment of orthologue families.

## Materials and methods

### Model optimization and data

We downloaded 14,526 coding sequence alignments for mammalian orthologue families and the corresponding inferred phylogenetic trees from the OrthoMaM (version 8) database [25]. These included sequences from 43 completely sequenced mammalian genomes, spanning from platypus to placental mammals. We restricted to the 12,336 families with at least 20 taxa for which the stop codon was included in the sequence alignment and positionally homologous with the end of the alignment. For each sequence alignment, we first re-estimated branch lengths of the phylogenetic trees using a the Muse and Gaut model [19] with the F1X4 model of codon equilibrium frequencies (MGF1X4), implemented in codonPhyml [45]. Treating the tree topology and relative branch lengths estimated by codonPhyml as fixed, we then optimized the model in equation 1 over the parameters *κ, ω, ϕ* and a scaling factor, *s*, by which we multiplied all branch lengths of the tree (in practice the scaling factor was typically close to 1 – interquartile range: 0.96 - 0.98). The model was implemented in R [23], and optimized using the optim function with the Nelder-Mead [24] method. Likelihood ratio tests were used to identify genes with evidence of conserved stop codons, with twice Δ*L* (the difference in the log likelihood of a model with *ϕ* fixed at 1 and a model with *ϕ* set to its maximum likelihood value) compared to a *χ*^2^ distribution with one degree of freedom. Code and data to reproduce our results are available from https://github.com/cseoighe/StopEvol.

### Simulations

We first produced simulated data under the model in equation 1 and optimized the parameters of the same model. Coding-sequence alignments for 1000 genes (and the corresponding phylogenetic trees) were sampled at random from the OrthoMaM database. We re-estimated the branch-lengths of the tree using a codon model (MGF1X4) implemented in codonPhyml [45]. Codons at the root of the tree were sampled from their equilibrium frequency under the F1X4 model (the F1X4 model sets codon frequencies to the product of the frequencies of their constituent nucleotides). Coding sequences were evolved from the root node over the phylogeny using code written in R. Model parameters were estimated from the simulated data as described above. We also simulated data under a Goldman and Yang [20] model (GY) with empirical target triplet frequencies derived from intron sequences of the same gene. Intron sequences from human protein-coding genes were derived from Ensembl Genes 94 [27]. We downloaded cDNA sequences and exon coordinates using Biomart and subtracted the exonic regions from the cDNA sequences. The first 100bp and last 100bp of each intron were discarded to reduce the impact of splicing signals on triplet frequencies. Codons were sampled from the intronic triplet frequencies for the root node of each tree and coding sequences were again simulated over the phylogeny.

### Mixture model, bootstrapping and simulation

We used a mixture model to estimate the proportion, *p*, of alignments for which the stop codon was under purifying selection pressure. Conditional on belonging to this set of alignments the value of *ϕ* was treated as a free parameter, while *ϕ* was equal to 1, otherwise. For tractability, we set *ω, kappa* and the tree scaling parameter to their maximum likelihood values. We performed a bootstrap over alignments to assess uncertainty in the estimates of *p* and *ϕ*. To test the accuracy with which the proportion of stop codons under purifying selection could be recovered we performed additional simulations. We simulated 1,000 genes with neutrally-evolving stop codons (*ϕ* = 1) and a further 1,000 genes with *ϕ* a normal random variable with mean 0.3 and standard deviation 0.1. Both sets of genes were simulated under a GY model with empirical triplet frequencies derived from intron sequences, as described above. We then performed 100 simulations. For each simulation we sampled 1000 genes from the two sets above, a random proportion (uniformly sampled from 0.1 to 0.8) of which were from the purifying selection set. We then applied our method (using the MGF1X4 model) to estimate the proportion of genes under purifying selection. Despite the substantial differences between simulation and the model the results are highly correlated (*R*^2^ = 0.996) and show only a very slight downward bias in the estimates (fig. S2). We also applied the mixture model using the GY model with the F3X4 model of codon frequencies, but found that this yielded less accurate results, despite being closer to the model under which the simulation was performed (fig. S6).

### Gene properties and enrichment analysis

Sequences of 3’ untranslated regions (UTRs) for human and mouse were downloaded from Ensembl Genes 91 [27]. Lengths of UTRs and coding regions were compared between genes that showed evidence of stop codon conservation (*ϕ* < 1 and *p* < 0.05) and the remaining genes using Wilcoxon rank sum tests. We performed tests of enrichment using DAVID (version 6.8) [28, 29].

### Expression level, expression breadth and mRNA half-life

Expression level and breadth were calculated using median values, per tissue, of gene transcripts per million (TPM) data from GTEx (version 7; [46]), downloaded on 8 February, 2018. We used the median of the per tissue median TPM values as a measure of expression level and the number of tissues in which the median TPM was greater than 10 as a measure of expression breadth. mRNA half-life data is from [47].

## Acknowledgments

We are grateful to Estienne Swart and Gary Loughran for comments on the manuscript. C.S. is supported by Science Foundation Ireland, award number 16/IA/4612. P.V.B. is supported by SFI-HRB-Wellcome Trust Biomedical Research Partnership [210692/Z/18/Z]. S.J.K. wishes to acknowledge personal support from the Irish Research Council.

## Author Contributions

CS initiated the project, developed the model, wrote the code, performed analysis and drafted the manuscript. PVB, HY and SK suggested and performed further analyses. AP developed and maintains the software repository.

## Additional Information

The authors declare that they have no competing interests.

## Supporting information

**S1 Fig.**
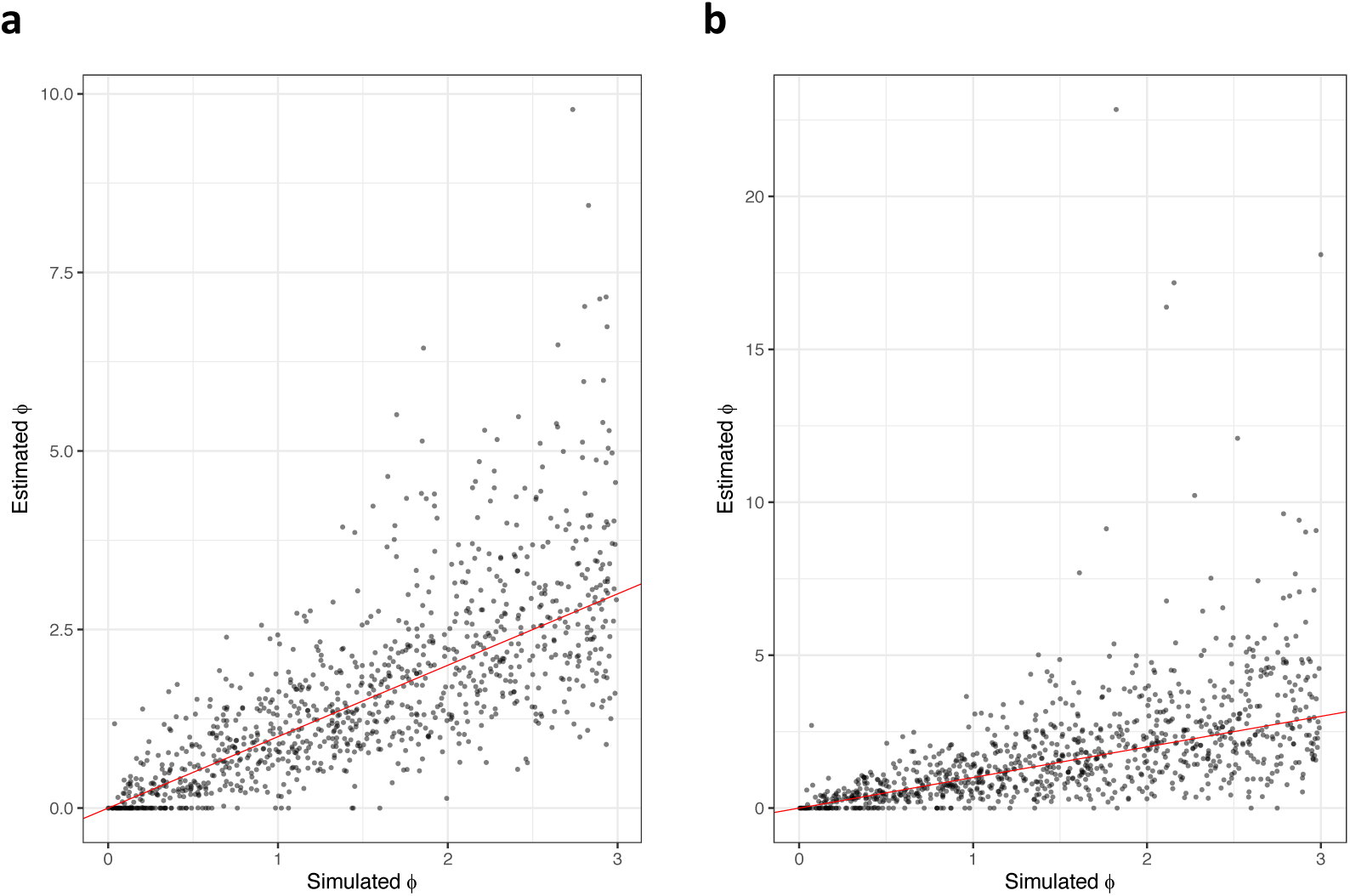
Simulation results for estimation of *ϕ*. Results of simulations based on 1,000 orthologue alignments, randomly sampled from OrthoMaM. (a) Sequences were simulated over the phylogenetic trees corresponding to the sequence alignments, under a stop-extended codon model based on MG with the F1×4 model of codon frequencies. Simulated values of *ϕ* were sampled uniformly between 0 and 3. Branch lengths of the phylogenies were re-estimated from the simulated alignments, under the MG F1×4 model, using codonPhyml. The stop-extended codon model was then fitted to the simulated alignments. The figure shows the maximum likelihood estimates, plotted against simulated values of *ϕ*. The identity line is shown in red. (b) Sequences were simulated over the phylogenetic trees corresponding to the sequence alignments, under a stop-extended codon model based on GY with empirical codon frequencies obtained from introns of the same genes. Only genes with at least 1000bp of introns were used (see Methods for details). Simulated values of *ϕ* were sampled uniformly between 0 and 3. Branch lengths of the phylogenies were re-estimated under a different model to that used for the simulation (MG F1×4), using codonPhyml. The stop-extended codon model, based also on MG F1×4 was then fitted to the simulated alignments. The figure shows the maximum likelihood estimates, plotted against simulated values of *ϕ*. The identity line is shown in red.

**S2 Fig.**
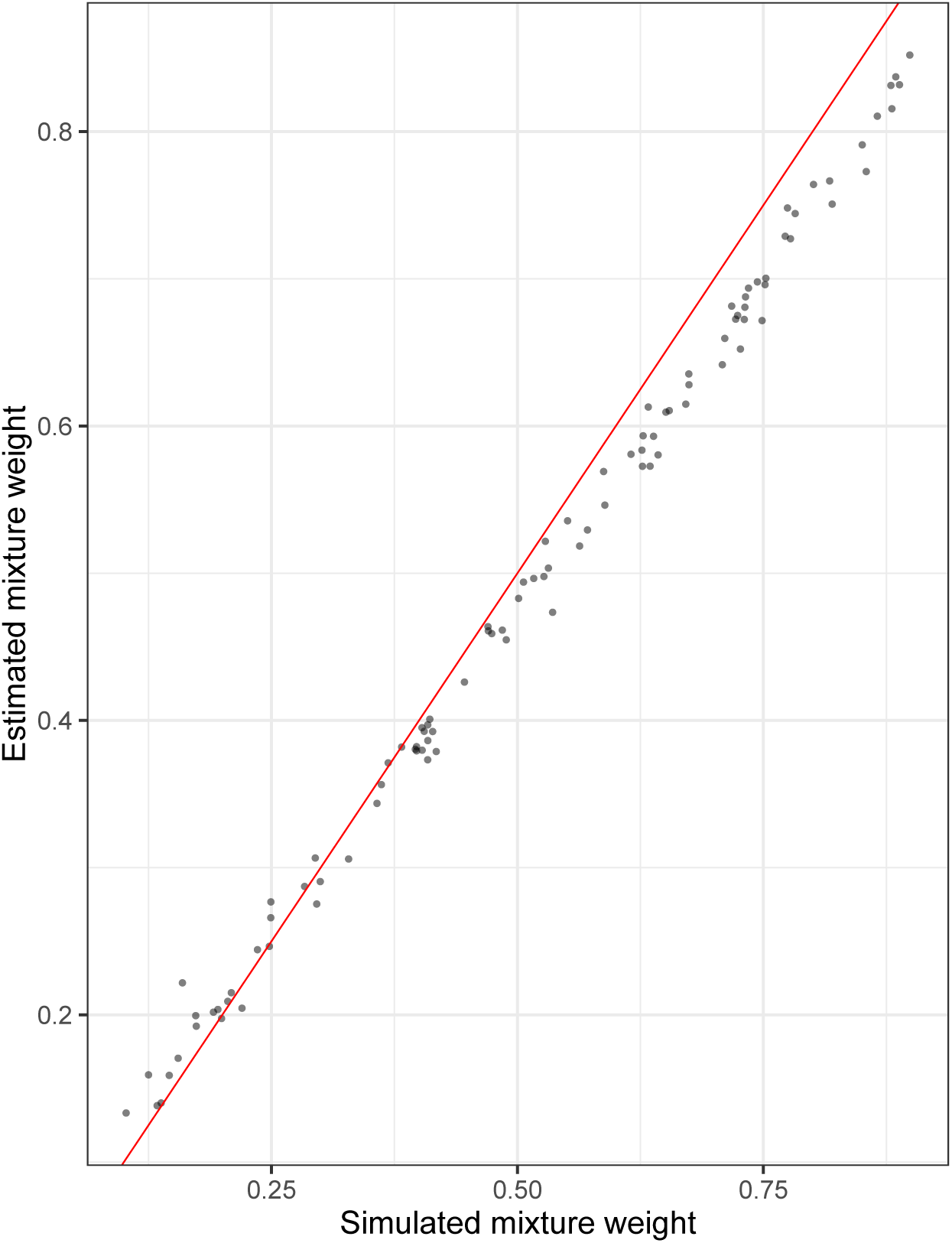
Simulation results for mixture model. Results of simulation to assess the accuracy of the estimate of the proportion of stop codons under selective constraint from the mixture model. Data was simulated under the GY model with empirical codon frequencies derived from intron sequences, as described in Methods. For each of 100 simulations, the proportion of genes under purifying selection was sampled uniformly between 0.1 and 0.8. The mixture model was fitted to the resulting alignments, under the MG model with the F1×4 model of codon frequencies. The figure shows the maximum likelihood value of the mixture weight corresponding to purifying selection, plotted against the proportion of alignments in the simulation for which the stop codon was under purifying selection. The identity line is shown in red.

**S3 Fig.**
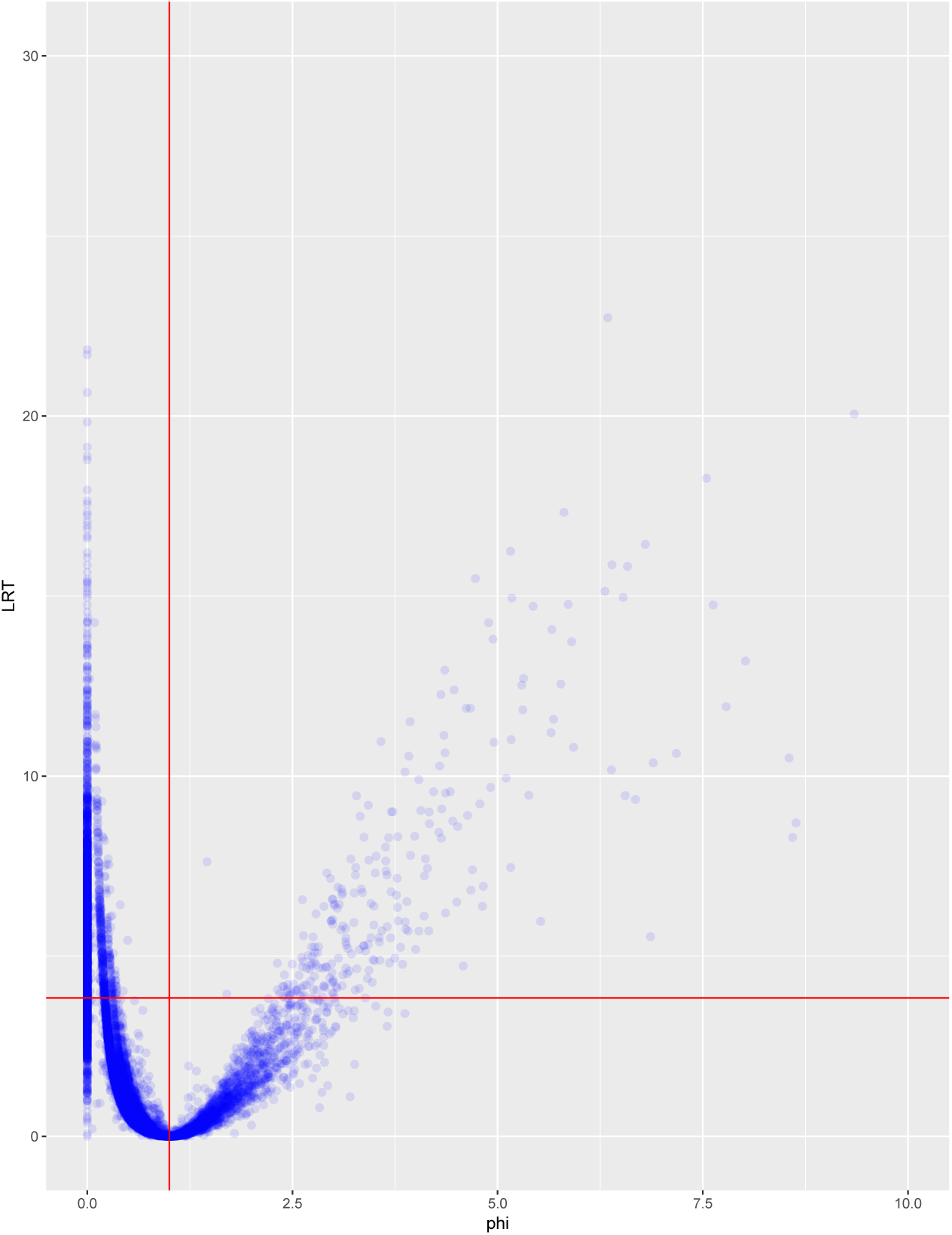
Results of model fit to individual orthologue alignments. Scatterplot showing the results of fitting the stop-extended model to the OrthoMaM alignments. The likelihood ratio test statistic (2Δ*lnL*) corresponding to the fit of the null model (*ϕ* = 1) compared to the alternative model (*ϕ* a free parameter) is plotted against the maximum likelihood estimate of *ϕ*. The horizontal line shows the critical value of the *χ*^2^ distribution with one degree of freedom and the vertical line shows *ϕ* = 1.

**S4 Fig.**
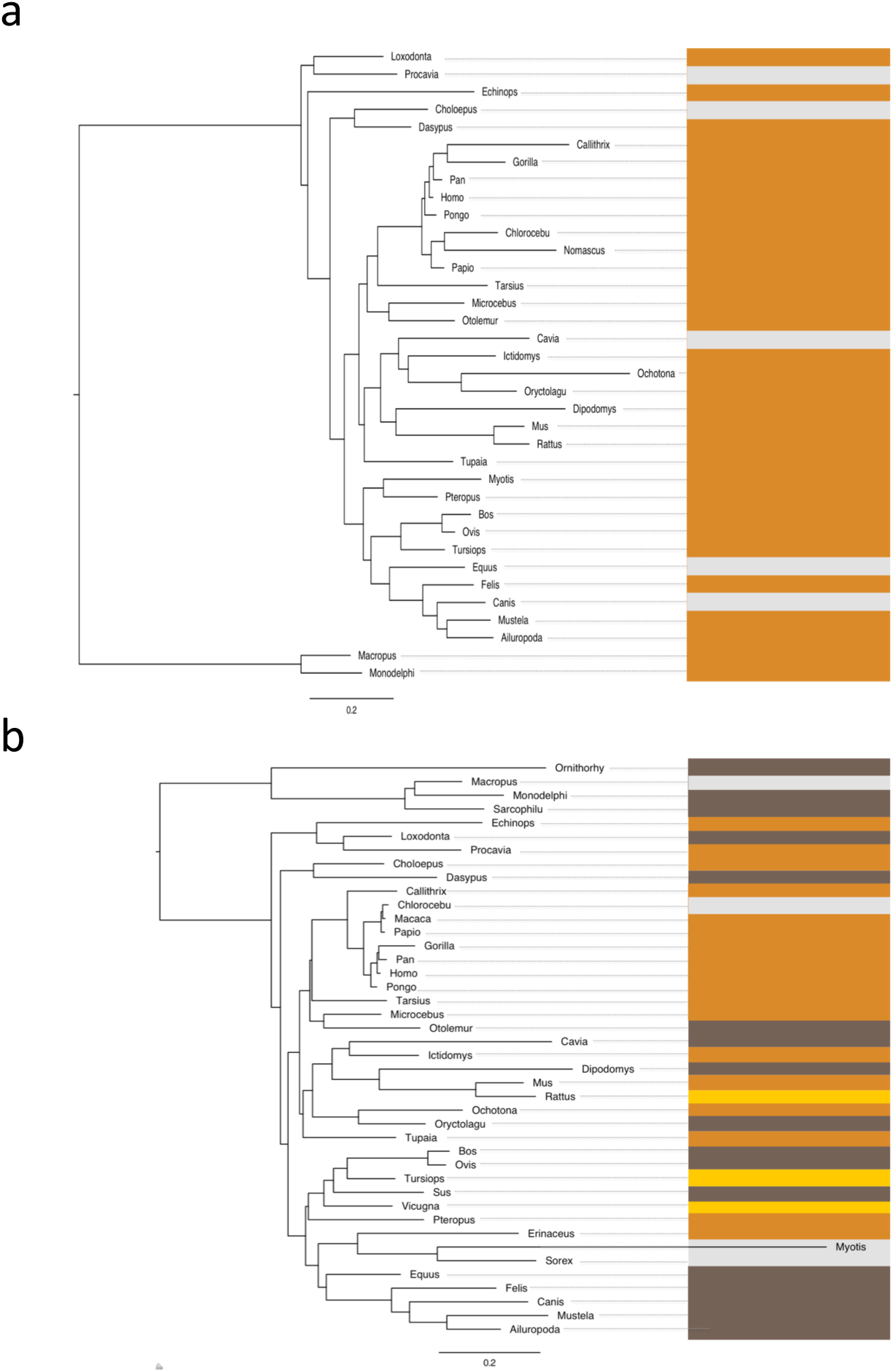
Example genes. An example of a gene with stop codon evolving at (a) a low and (b) a high rate. The phylogeny is coloured according to the stop codon colour nomenclature as in figure 1. Taxa with gaps in the sequence alignment at the stop codon are indicated in grey. The genes corresponding to panels a and b are *INSR* and *PARP1*, respectively.

**S5 Fig.**
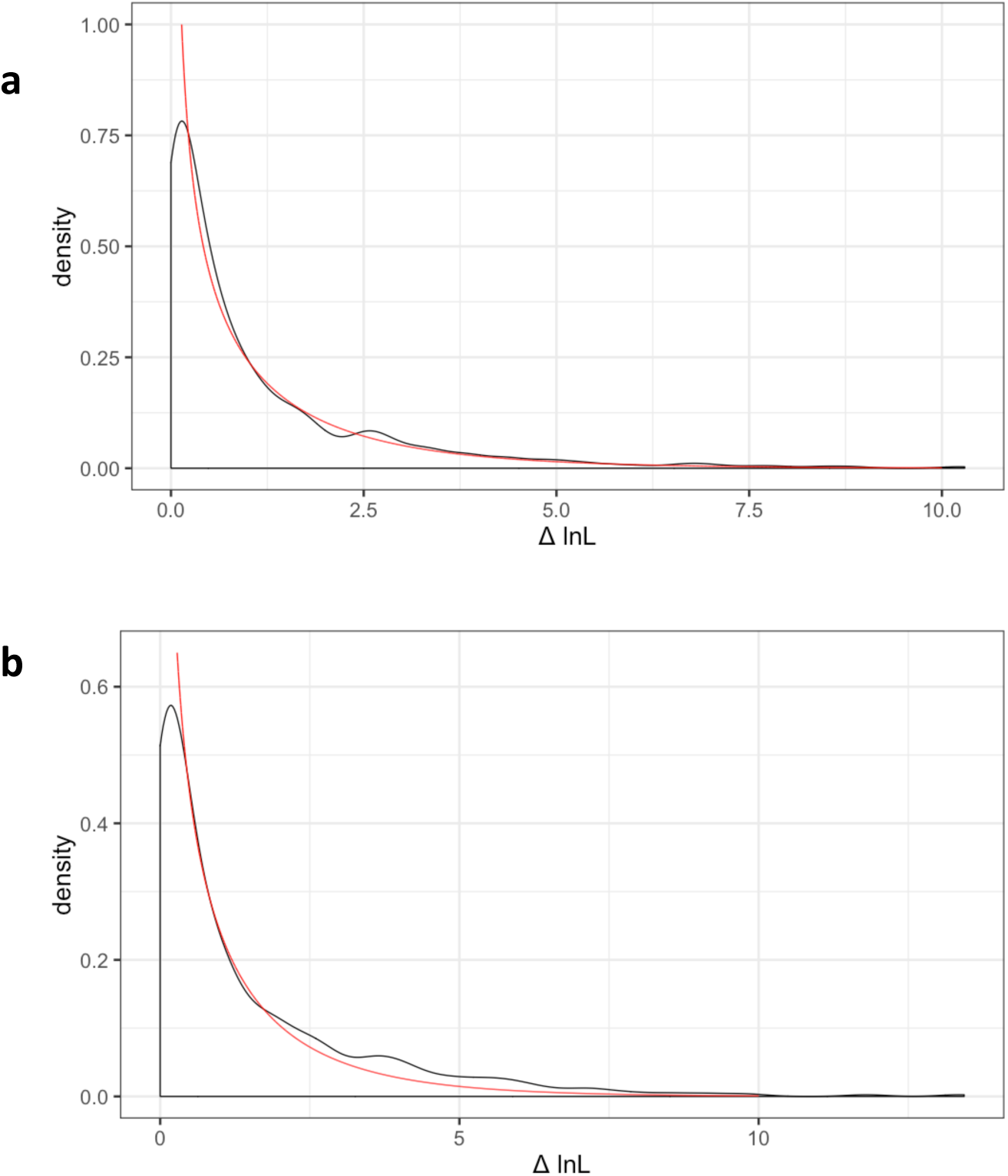
Simulation results for model comparison. Density plots of the likelihood ratio test statistic from the simulations shown in figure S1. The red lines show the *χ*^2^ distribution with one degree of freedom. a) simulations under the model b) simulations under an alternative model. See the legend of figure S1 for details of the models.

**S6 Fig.**
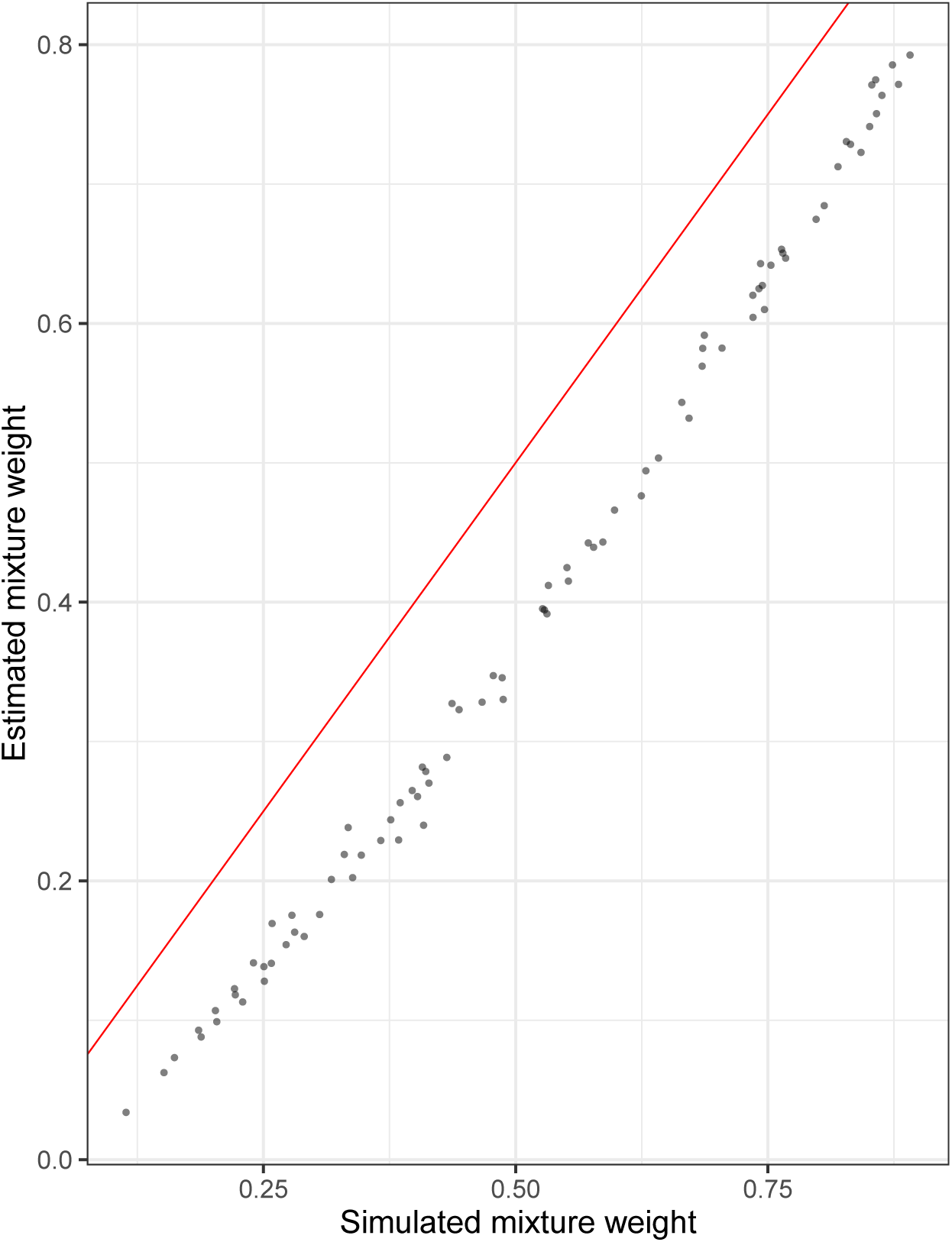
Model comparison simulations with different model forms. The same simulations as in figure S2 but in this case the GY model with the F3×4 model of codon frequencies was used.

